# *FAAH*, *SLC6A4*, and *BDNF* variants are not associated with psychosocial stress and mental health outcomes in a population of Syrian refugee youth

**DOI:** 10.1101/685636

**Authors:** Christopher J. Clukay, Anthony Matarazzo, Rana Dajani, Kristin Hadfield, Catherine Panter-Brick, Connie J. Mulligan

## Abstract

The developmental origins of health and disease (DOHaD) hypothesis posits that early childhood stressors disproportionately impact adult health. Numerous studies have found adult mental health to be associated with childhood adversities and genetic variants, particularly in genes related to neurochemistry. However, few studies have examined the way interactive effects may manifest over time and fewer still include protective factors, like resilience. Our group has previously found associations between the monoamine oxidase A gene, *MAOA*, and a contextually-specific measure of resilience with a measure of perceived psychosocial stress over time in Syrian refugee youth. In this study, we work with the same sample of adolescents to test genetic variants in three additional candidate genes (*FAAH*, the 5-HTTLPR region of *SLC6A4*, and *BDNF*) for associations with six psychosocial stress and mental health outcomes. Using multi-level modeling, we find no association between variants in these candidate genes and psychosocial stress or mental health outcomes. Our analysis included tests for both direct genetic effects and interactions with lifetime trauma and resilience. Negative results, such as the lack of genetic associations with outcome measures, provides a more complete framework in which to better understand positive results and associations.

## Introduction

Early life stressors have long been noted to have a disproportionate impact on adult health (Barker et al., 1993). A well-known example of this phenomenon is the Dutch Hunger Winter Families Study, which has documented health effects in the adult offspring of pregnant mothers exposed to starvation during the winter of 1944-1945 at the end of World War II (Lumey et al., 1993). Maternal exposure to starvation while pregnant was associated with multiple phenotypes in adult offspring, including increased blood pressure, glucose intolerance, increased BMI, and worsened mental health (Lumey et al., 2007; Roseboom et al., 2006; Stein et al., 2009; Tobi et al., 2018; Yarde et al., 2013). Effects were also seen from post-natal exposure, with girls who were exposed between birth and age nine displaying increased risk for obesity in adulthood (van Abeelen et al., 2012). These long-term effects highlight the importance of understanding early life adversity and the factors that may modify the impact of adversity. Ongoing civil wars and related humanitarian crises highlight the need to better understand the multiple effects of early life adversity.

Accordingly, our research team has studied the impact of exposure to traumatic events and the factors underlying associations with psychosocial stress and mental health outcomes in Syrian refugee youth. Using a longitudinal method over the course of one year, our team identified factors related to genetics, a contextually-specific measure of resilience, and a psychosocial intervention (Clukay et al., in press; Panter-Brick et al., 2018a; Panter-Brick et al., 2017). Specifically, we found associations between *MAOA* genetic variants and levels of resilience with perceived stress in males in our study population (Clukay et al., in press). Here, we expand upon our previous genetic study to investigate genetic variants in three other genes of interest (*FAAH*, the 5-HTTLPR region of *SLC6A4*, and *BDNF*).

The fatty acid amide hydrolase gene (*FAAH*) encodes an enzyme by the same name (FAAH) that degrades ethnocannabinoids in the nervous system, which helps to regulate anxiety and mood among other functions. Altered FAAH enzyme activity has been directly linked to neurological changes that increase excitability and anxiety (Gunduz-Cinar et al., 2013). Furthermore, the specific genetic variant of *FAAH* assayed in our study (rs324420) has been previously associated with several anxiety disorders, particularly in association with psychosocial stress (Lazary et al., 2016; Spagnolo et al., 2016).

The serotonin transporter gene (*SLC6A4*, or solute carrier family 6 member 4) encodes a transport protein which removes serotonin from the synaptic cleft. Removal of serotonin means that the neurotransmitter is essentially deactivated, which is why SLC6A4 is a common target for antidepressant medications (Wilkie et al., 2008). The serotonin-transporter-linked polymorphic region (5-HTTLPR) in the promoter of *SLC6A4* has over half a dozen repeat polymorphisms that affect transcription of the gene (Murdoch et al., 2013). Variation in the 5-HTTLPR region has been linked to multiple psychosocial stress and mental health phenotypes, including obsessive-compulsive disorder and, in combination with childhood trauma, memory bias and riskier decision making (Hu et al., 2006; Stoltenberg et al., 2011; Vrijsen et al., 2015).

Finally, the brain-derived neurotrophic factor gene (*BDNF*) encodes an enzyme which acts on a wide variety of neurological systems, including regulation of neural growth and plasticity, as well as interaction with dopamine (Kowiański et al., 2018). The variant assayed in this study (rs6265) results in a valine to methionine substitution. The methionine allele results in altered phosphorylation and decreased enzyme in the synaptic cleft by interfering with intracellular transport (Deng et al., 2013; Hempstead, 2015). These altered transport patterns result in changes to the hippocampus, which have been linked to depression and other phenotypes, particularly in combination with altered cortisol and early life stress (Frodl et al., 2014; Herbert et al., 2012). The variant has also been linked to human resilience, when defined as the individual capacity to cope with stress and adversity (Kang et al., 2013). Kang et al. found that *BDNF* genotype interacted with other genetic factors in association with the Connor-Davidson Resilience Scale, specifically in males, indicating possible sex-specific effects.

In this study, we test for associations between three candidate genes, lifetime trauma exposure, and resilience levels with six measures of psychosocial stress and mental health in a longitudinal sample of Syrian refugee youth who re-settled in Jordan. We use multi-level modeling to test genetic variants for direct associations with psychosocial stress and mental health outcomes, as well as interactive effects with trauma exposures and resilience levels.

## Materials and Methods

### Study design

The design for this study has been detailed in previous publications (Clukay et al., in press; Dajani et al., 2018; Panter-Brick et al., 2019). In brief, buccal samples and survey data were collected from Syrian refugee youth (ages 11-18), forcibly displaced to Jordan. Participants resided in four sites in northern Jordan (Irbid, Jarash, Mafraq, and Zarqa), and were enrolled in a psychosocial intervention (*Advancing Adolescents*, an 8-week program of structured activities) delivered by Mercy Corps to children and adolescents affected by the Syria crisis. Buccal swabs and survey data on trauma, resilience, psychosocial stress, and mental health were collected at Week 1, ~Week 13, and ~Week 48. Samples and data were collected in two cohorts that were combined for analysis; March-June 2015 (*n*=103) and September 2015-February 2016 (*n*=297).

This study was approved by the Prime Minister’s Office of Jordan and the Yale University IRB (ID 1502015359). Informed consent was obtained in Arabic from both the participating adolescents and their parent or guardian. All participants agreed to provide a buccal sample. All data analyzed in this study are available on Mendeley Data (doi:10.17632/2wbptg7vyn.3).

### Trauma, resilience, psychosocial stress and mental health measures

Lifetime trauma exposure was measured using a modified version of the Traumatic Events Checklist, adapted for use in this region and age group (Panter-Brick et al., 2009). Resilience was measured using the Child and Youth Resilience Measure-12 (CYRM-12), a self-reported scale specifically designed to assess strengths for youth living in adversity (Panter-Brick et al., 2017; Ungar and Liebenberg, 2011). Resilience is defined using a socioecological paradigm and measures protective factors at individual, relational, and contextual levels. Since the CYRM-12 measure was developed during the project, resilience measures were only available for the second cohort (*n* = 163 males and 127 females, for a total of 290 participants).

Six psychosocial stress and mental health outcomes were collected for this study. The Perceived Stress Scale (PSS) measures the degree to which an individual views situations in life as stressful (Cohen et al., 1983). The version of the PSS used here has been translated into Arabic and validated for use in Jordan (Almadi et al., 2012). The Human Distress Scale (HD) was developed for use in conflict-affected populations in the Middle East region to assess feelings of anxiety regarding lack of control over an individual’s life (Hammoudeh et al., 2013). The Human Insecurity Scale (HI) was developed alongside HD to measure individual fears regarding threats to themselves and to their family, community, and future (Hammoudeh et al., 2013; Ziadni et al., 2011). The Arab Youth Mental Health Scale (AYMH) screens for anxiety and symptoms of depression, and was specifically designed for use in Middle Eastern populations (Mahfoud et al., 2011; Makhoul et al., 2011). The Strengths & Difficulties Questionnaire (SDQ) measures emotional and peer-relation problems (Alyahri and Goodman, 2006; Goodman and Goodman, 2009). The Children’s Revised Impact of Events Scale (CRIES-8) assesses symptoms of post-traumatic stress after specific trauma exposure (Punamäki et al., 2015; Veronese and Pepe, 2013).

### Swab collection and DNA extraction

Cheek swabs were collected using either DNA Buccal Swabs (Isohelix, United Kingdom) or Transport Swabs (APCO Laboratory Consumable Plastic, Jordan). Before giving a sample, all participants rinsed their mouths and brushed the swab against their cheek for up to 30 seconds. Depending on swab type, DNA extractions were performed using either the Xtreme DNA Isolation Kit (for DNA Buccal Swabs; Isohelix, United Kingdom) or the Qiagen DNA Investigator Kit (for Transport Swabs; Qiagen, USA). DNA extractions were performed as recommended by the manufacturer except that the AW2 wash was performed twice for swabs extracted using the Qiagen kit in order to increase DNA purity.

### Genotyping Assays and Allele Coding

#### FAAH

Protocols for assaying rs324420 in *FAAH* were based on a modification of a previous study by Monteleone et al. (2009). Samples were amplified using the following primers: Forward 5′-ATG-TTG-CTG-GTT-ACC-CCT-CTC-C-3′, Reverse 5′-CAG-GGA-CGC-CAT-AGA-GCT-G-3’. The reaction consisted of 1 µL of genomic DNA, 12.5 µL of 2x Platinum™ Hot Start PCR Master Mix (Invitrogen, USA), 0.25 µL of 10 µM forward primer, 0.25 µL of 10 µM reverse primer, 0.75 µL dimethyl sulfoxide, and 10.25 µL water for a total volume of 25 µL. The cycling profile was as follows: 95°C for 5 min followed by 34 cycles of 94°C for 30 sec, 62°C for 30 sec, and 72°C for 30 sec, followed by a final three-minute step at 72°C.

Following amplification, the product was digested for one hour at 37°C using 40 U *Eco*O109I (New England Biolabs, USA), 5 µL accompanying buffer solution, and water for a final reaction volume of 30 uL. All digested products were electrophoresed on 3% agarose gels using Agarose SFR™ (VWR Life Science, USA) for 3 hours at 80V. Individuals were coded based on the presence or absence of the minor allele (i.e. Pro/Pro vs. Pro/Thr + Thr/Thr), consistent with previous findings in this gene (Bühler et al., 2014; Lazary et al., 2016; Spagnolo et al., 2016).

#### 5-HTTLPR

A repeat polymorphism and associated single nucleotide variant (rs25531) in the 5-HTTLPR of *SLC6A4* were assayed. Four categories of the repeat polymorphism (XS, S, L, and XL) were assayed based on a modification of a published protocol by Caspi et al. (2003) using the GoTaq® PCR Core System I (Promega, USA). The following primer sequences were used: Forward 5’-ATG-CCA-GCA-CCT-AAC-CCC-TAA-TGT-3’, Reverse 5’-GGA-CCG-CAA-GGT-GGG-CGG-GA-3’. The reaction consisted of 3 µL genomic DNA, 2.5 µL of 25 mM MgCl_2_, 10 µL of 5X Green GoTaq® Flexi Buffer, 0.25 µL GoTaq Taq DNA polymerase, 1 µL of 10 mM nucleotide mix, 0.6 µL of 10 µM forward primer, 0.6 µL of 10 µM reverse primer, and 32.05 µL water for a total volume of 50 µL. This MgCl_2_ concentration is lower than in many standard master mixes, consistent with previous literature regarding this reaction (Yonan et al., 2006). The cycling profile was as follows: 95°C for 5 min followed by 34 cycles of 94°C for 30 sec, 66°C for 30 sec, and 72°C for 30 sec, followed by a final three-minute step at 72°C.

An A/G polymorphism (rs25531) present in L variants was assayed in all amplification products by digestion with 24U of *Msp*I (New England Biolabs) overnight at 37°C as recommended by Wang et al. (2011). Digested products were electrophoresed on 3% agarose gels using Agarose SFR™ (VWR Life Science, USA) for 3 hours at 80V, followed by a 20-minute immersion in dilute ethidium bromide before imaging.

Combined repeat and A/G polymorphisms were separated into two categories of high and low expression variants based on the literature (Iurescia et al., 2016). All XS and S variants (≤ 14 repeats), as well as all L variants (16 repeats) containing a G at rs25531, were coded as low expression (Ehli et al., 2011; Wang et al., 2011). All L variants containing an A at rs25531 were coded as high expression. Due to the complex nature of the XL alleles (> 16 repeats), whose expression has rarely been studied and can vary depending on which segments were duplicated, the four individuals exhibiting these variants were excluded from the analysis. A dosage scheme was used to code individuals based on genotype. Individuals with two low expression variants were assigned a score of zero, individuals with one low expression variant and one high expression variant were given a score of 0.5, and individuals with two high expression variants were assigned a score of one.

#### BDNF

Protocols for assaying rs6265 in *BDNF* were based on a modification of previous work by Miller et al. (2013). Samples were amplified using the following primers: Forward 5′-ATC-CGA-GGA-CAA-GGT-GGC-3′, Reverse 5′-CCT-CAT-GGA-CAT-GTT-TGC-AG-3′. The reaction consisted of 1 µL genomic DNA, 15 µL of 2x GoTaq Hot Start Green Master Mix (Promega, USA), 0.3 µL of 10 µM forward primer, 0.3 µL of 10 µM reverse primer, and 13.4 µL water in a final reaction volume of 30 µL. The cycling profile was as follows: 95°C for 5 min followed by 34 cycles of 94°C for 30 sec, 60°C for 30 sec, and 72°C for 30 sec, followed by a final three-minute step at 72°C.

The product was digested overnight at 37°C using 20 U *Pml*I (New England Biolabs, USA), 5 uL accompanying buffer, and water for a final reaction volume of 50 uL. All digested products were electrophoresed on 2% agarose gels using Agarose SFR™ (VWR Life Science, USA) for 2 hours at 80V. Individuals were coded based on the presence or absence of the minor allele (i.e. Val/Val vs. Val/Met + Met/Met) as is standard for this variant (Frodl et al., 2014; Gerritsen et al., 2012; Kang et al., 2013).

### Statistical Analyses

All statistical analyses were conducted in R (R Development Core Team, 2015). Multi-level modeling was used to test for the significance of all associations using the nlme package (Pinheiro et al., 2016). Modeling structure is based on Clukay et al. (in press). A nested model structure was used in which timepoints were grouped within individuals. Significance tests for variables of interest were conducted by performing an ANOVA on nested models, one with the variable of interest present and one without it. Both age and collection site were included as time-invariant, fixed effect covariates. An 8-week stress attunement intervention program (*Advancing Adolescents*) was administered to the study population between baseline and the first follow-up visit. This intervention was included as a time-varying covariate where all individuals were coded as 0 at baseline and those who underwent the intervention were coded as 1 for all follow-ups. Three sets of models were run; one testing for direct genetic effects, one testing for an interaction with trauma, and one testing an interaction with resilience in which trauma was a covariate. Consistent with our previous studies (Clukay et al.; Dajani et al., 2018; Panter-Brick et al., 2018a), trauma was treated as time-invariant; trauma exposure data were collected at baseline. Resilience was treated as a time-varying measure, meaning the value at each timepoint was included. All model coefficients and structures are detailed in Supplemental Tables S1-S4.

Time-invariant terms have both intercept effects and slope effects. Intercept effects are associated with an individual’s score for an outcome measure, but the effect is independent of time. In contrast, slope effects are associated with an individual’s trajectory of the outcome over time and, thus, time is included in the term.

In order to test whether sufficient variation existed to use multi-level modeling, we examined whether a significant association existed between time and the outcome variables. When testing individuals assayed for *FAAH*, five of the six outcomes (PSS, HD, AYMH, SDQ, and CRIES-8) had sufficient variation with time (*p* < 0.05). This finding remained true when testing the subset of individuals with resilience measures. When testing individuals assayed for 5-HTTLPR, four of the outcomes had sufficient variation over time (PSS, HD, AYMH, SDQ). When testing the subset of individuals with resilience measures, these four outcomes plus CRIES-8 has sufficient variation over time. A number of studies have noted sex-specific associations with *BDNF* genotype (Herbert et al., 2012; Kang et al., 2013). Thus, the study sample was subdivided into males and females to test for associations between our outcomes and *BDNF*. Five outcomes had sufficient variation to test for association in females (PSS, HD, AYMH, SDQ, and CRIES-8), but only four outcomes had sufficient variation in males (PSS, HD, AYMH, and SDQ). When testing the subset of individuals with resilience measures, the same outcomes had sufficient variation in females but only three outcomes had sufficient variation in males (PSS, HD, and SDQ). HI lacked sufficient variation to test for association with any of the three genetic variants in any model.

The multiple testing significance threshold included correction for the five psychosocial stress and mental health outcomes with sufficient variation for modeling, two sexes, and three types of possible associations (direct association, interaction with trauma, and interaction with resilience). This resulted in a significance threshold of *p* = 0.05 / (5 × 2 × 3) = 1.6 × 10^−3^.

## Results

### Sample Characteristics

Sample characteristics for the study population can be found in Table 1. Regarding the three genetic variants, none exhibited a significant difference in allele frequencies between males and females (*p* > 0.1 in all cases). Males reported more traumatic events than females, though comparable levels of resilience were seen in both groups. Genotype frequencies for all three genes did not vary significantly by either levels of trauma or resilience. Regarding the outcome measures, females demonstrated higher scores on the PSS, HD, AYMH, and SDQ measures, signifying increased psychosocial stress and worsened mental health at baseline, compared to males.

**Table 1.** Summary of sample characteristics at baseline.

|  |  | Males | Females |
| --- | --- | --- | --- |
| Participants (#) |  | 221 | 179 |
| Individuals with at least one other timepoint (#) |  | 137 | 126 |
| Age (years) |  | 14.2<br>(1.81) | 14.5<br>(1.90) |
| *BDNF* | Val/Val (#) | 146 | 108 |
|  | Val/Met (#) | 61 | 61 |
|  | Met/Met (#) | 11 | 7 |
| *FAAH* | Pro/Pro (#) | 136 | 121 |
|  | Pro/Thr (#) | 73 | 47 |
|  | Thr/Thr (#) | 7 | 7 |
| **5-HTTLPR**<br>**region of**<br>***SLC6A4*** | **High Expression (#)** | 50 | 48 |
|  | **High-Low**<br>**Expression**<br>**Heterozygotes (#)** | 104 | 72 |
|  | **Low Expression (#)** | 61 | 53 |
| **Lifetime Traumatic Events\*** |  | 6.77<br>(3.26) | 5.86<br>(3.22) |
| **Child and Youth Resilience**<br>**Measure (CYRM-12)** |  | 49.8<br>(6.72) | 49.1<br>(7.06) |
| **Perceived Stress Scale\* (PSS)** |  | 28.2<br>(3.12) | 29.4<br>(6.44) |
| **Human Distress Index\* (HD)** |  | 39.9<br>(19.2) | 44.7<br>(22.9) |
| **Human Insecurity Index (HI)** |  | 66.3<br>(20.0) | 70.1<br>(19.3) |
| **Arab Youth Mental Health\***<br>**(AYMH)** |  | 34.7<br>(7.68) | 37.3<br>(9.25) |
| **Strengths & Difficulties**<br>**Questionnaire\* (SDQ)** |  | 15.0<br>(5.58) | 16.3<br>(6.23) |
| **Children's Revised Impact of**<br>**Events Scale (CRIES-8)** |  | 19.7<br>(10.6) | 19.3<br>(11.6) |

Unless otherwise stated, values are mean (SD) at baseline. Alleles for the *BDNF* gene were classified based on whether they code for valine (Val) or methionine (Met) amino acids. Alleles for the *FAAH* gene were classified based on whether they code for proline (Pro) or threonine (Thr) amino acids. For 5-HTTLPR, L variants with an A at rs25531 were classified as high expression, and XS and S variants plus L variants with a G at rs25531 were classified as low expression. An asterisk (\*) denotes a significant difference between males and females at a p-value threshold of 0.05. The resilience measure was only collected for a subset of 163 males and 127 females.

### Association of Genetic Variants and Outcome Measures

We analyzed both males and females together when testing *FAAH* and 5-HTTLPR. *BDNF* has been previously noted to exhibit sex-specific associations (Kang et al., 2013), so we tested each sex separately for analysis of *BDNF*, though results were similar when the sexes were combined (data not shown). After multiple testing correction, no significant associations were seen between any of the outcome measures and *FAAH* or 5-HTTLPR in the complete dataset, or *BDNF* in males or females (see Fig 1 where darker shades of red denote smaller p-values). Time-invariant terms in Models 1 and 2 have both intercept and slope terms, e.g. ‘*FAAH*’ and ‘*FAAH* x Time’, respectively (see Methods for details on this distinction). Neither direct genetic effects nor interactions of genetic effects with either trauma or resilience were significant for any of the outcomes measures after correction for multiple testing.

**Figure 1.**
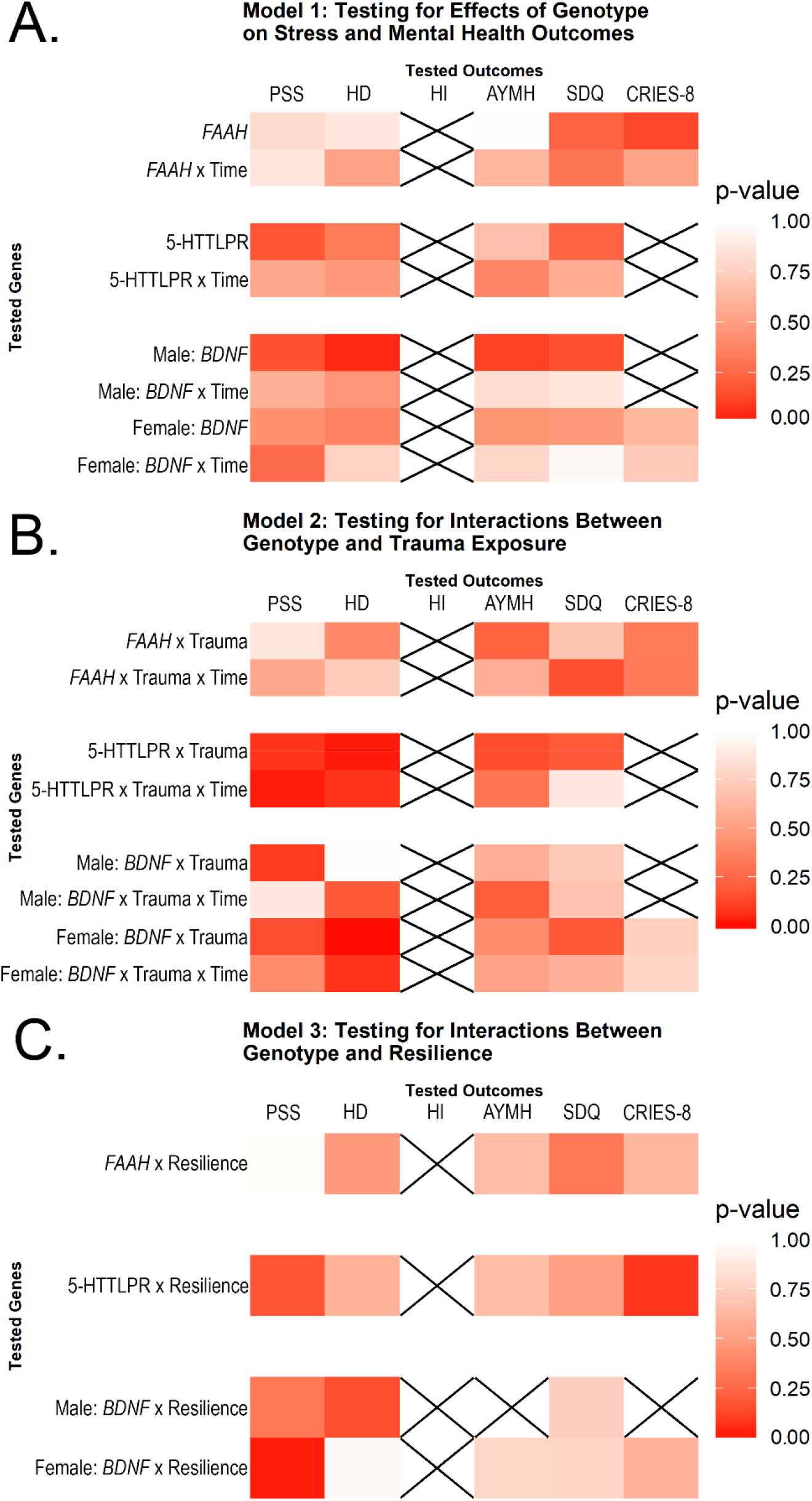
Heatmap of potential associations between tested genes and outcome measures. Model 1 examined direct effects of the tested gene and Models 2 & 3 examined interactive effects between the gene and trauma or resilience, respectively. Crossed-out squares indicate there was insufficient variance in that measure over time to test using multi-level modeling. Darker shades of red indicate smaller p-values. None of the reported results met the multiple testing threshold of 1.6 × 10^−3^. Abbreviations are as follows: PSS = Perceived Stress Scale, HD = Human Distress Scale, HI = Human Insecurity Scale, AYMH = Arab Youth Mental Health Scale, SDQ = Strengths & Difficulties Questionnaire, CRIES-8 = Children’s Revised Impact of Events Scale. Note there is no slope term for the resilience models because resilience was treated as a time-varying measure, meaning variance over time is inherently included.

Both Perceived Stress Scale (PSS) and Human Distress (HD) exhibited several potential associations, but these did not survive multiple testing correction (*p* ≤ 1.6 × 10^−3^). In relation to PSS, a possible interaction between 5-HTTLPR and Trauma over time was noted (‘5-HTTLPR x Trauma x Time’; *p* = 0.028) as was an interaction between *BDNF* and resilience in females (‘*BDNF* x Resilience’; *p* = 0.027). Regarding HD, an interaction between 5-HTTLPR and Trauma was noted at intercept (‘5-HTTLPR x Trauma’; *p* = 0.025) as well as two associations with *BDNF* at intercept; a direct effect in males (‘*BDNF*’; *p* = 0.047) and an interaction with trauma in females (‘*BDNF* x Trauma’; *p* = 0.0098). The *BDNF* interaction with trauma in females is noteworthy in that it is the nearest to significance in the dataset. The other testable outcomes (AYMH, SDQ, and CRIES-8) and the other gene (*FAAH*) exhibited no statistically significant findings, even before correction for multiple testing (all *p* > 0.05). For details regarding model coefficients and standard errors, see Supplemental Tables S1-S4.

## Discussion

We use a longitudinal design with measures of psychosocial stress and mental health across the span of one year. Including multiple timepoints enables the use of multi-level modeling, which affords us greater statistical power than a traditional cross-sectional study as well as the ability to examine predictors of change over time. Our previous identification of significant associations between *MAOA* variants and one of the studied psychosocial stress outcomes (PSS) suggests that our study has sufficient power to detect such associations (Clukay et al., in press).

*FAAH* has been previously associated with psychosocial stress and anxiety, for which we have multiple measures including generalized scales developed for use globally (PSS) and specialized scales designed for use in the region (HD and AYMH; Spagnolo et al., 2016). However, few studies have linked *FAAH* specifically with childhood trauma, Lazary et al. (2016) being the notable exception that led us to assay the gene. Thus, considering the paucity of support in the literature and the fact that *FAAH* was the only gene in which no p-value approached significance (all p-values > 0.05), it is possible that *FAAH* may simply not be a strong candidate to explore in relation to childhood adversity.

Both 5-HTTLPR and *BDNF* are traditionally associated with depressive symptoms (Dalton et al., 2014; Dwivedi, 2009). However, the only outcome measure in our study which partially captures depressive symptoms is AYMH, which has been noted to have only moderate sensitivity in girls and poor sensitivity in boys (Mahfoud et al., 2011). In addition, a recent study by Border et al. (2019) with over 115,00 participants assayed these genes as well as all variants commonly tested for association with depression (i.e. tested in more than 10 studies) and found no associations between genetic variation and any depressive phenotypes. Considering these factors, our results regarding a lack of association between either 5-HTTLPR or *BDNF* and the measured outcomes are not entirely unexpected.

Lastly, it is noteworthy that all near-significant associations (i.e. significant before multiple testing correction) were seen in either PSS or HD. These results are consistent with our previous findings regarding *MAOA*, in which we found an association with PSS after multiple testing correction. Furthermore, the fact that HD was developed specifically for use in conflict-affected populations in the Middle East may lend it increased power in our population. These findings may indicate that these outcome measures are worth pursuing with a larger sample.

Not all genes previously associated with psychosocial stress and mental health may have an effect in our study population, nor act via the same mechanism, especially given well-known differences between the short- and long-term effects of stress. Knowing which variables have been found *not* to associate with psychosocial stress and mental health over time is important for understanding the underlying mechanisms of the variables which are found to influence health outcomes. Further investigations into the genetics of psychosocial stress and mental health, particularly with respect to trauma and resilience over time, are needed. As previously noted by this team (Panter-Brick et al., 2018b), increased partnerships with both local populations and international aid groups are needed to ensure more culturally-sensitive studies and more effective assistance to vulnerable populations.

## Acknowledgments

Laboratory research was funded by the University of Florida College of Liberal Arts and Sciences (CJM). Field research was funded by Elrha’s Research for Health in Humanitarian Crises (R2HC) Programme (https://www.elrha.org/project/yale-psychosocial-call2/) which aims to improve health outcomes by strengthening the evidence base for public health interventions in humanitarian crises (https://www.elrha.org/r2hc) (CPB). The R2HC programme is funded equally by the Wellcome Trust and the UK Government. Additional funding from NSF grant DGE-1315138 (CJC). We gratefully acknowledge the Syrian families who participated in this project. We thank Dima Hamadmad, Ghufran Abudayyeh, Rahmeh Hyari, and Sana’a Bakeer from the Taghyeer Foundation for excellent fieldwork.

## Data Availability

All sample data are available on Mendeley Data (doi:10.17632/2wbptg7vyn.3). All models used in analysis are available in tables S1-S4.

